# Systematic Identification of Novel Cancer Genes through Analysis of Deep shRNA Perturbation Screens

**DOI:** 10.1101/807248

**Authors:** Hesam Montazeri, Mairene Coto-Llerena, Gaia Bianco, Ehsan Zangene, Stephanie Taha-Mehlitz, Viola Paradiso, Sumana Srivatsa, Antoine de Weck, Guglielmo Roma, Manuela Lanzafame, Martin Bolli, Niko Beerenwinkel, Markus von Flüe, Luigi M. Terracciano, Salvatore Piscuoglio, Charlotte K. Y. Ng

**Author notes:** equal author contribution (Hesam Montazeri, Mairene Coto-Llerena, Gaia Bianco). Hesam Montazeri Mairene Coto-Llerena Gaia Bianco Ehsan Zangene Stephanie Taha-Mehlitz Viola Paradiso Sumana Srivatsa Antoine de Weck Guglielmo Roma Manuela Lanzafame Martin Bolli Niko Beerenwinkel Markus von Flüe Luigi M. Terracciano Salvatore Piscuoglio Charlotte K. Y. Ng.

## Abstract

**Background:** Systematic perturbation screens provide comprehensive resources for the elucidation of cancer driver genes, including rarely mutated genes that are missed by approaches focused on frequently mutated genes and driver genes for which the basis for oncogenicity is non-genetic. The perturbation of many genes in relatively few cell lines in such functional screens necessitates the development of specialized computational tools with sufficient statistical power.

**Results:** Here we developed APSiC (<u>*A*</u>nalysis of <u>*P*</u>erturbation <u>*S*</u>creens for *i*dentifying novel <u>*C*</u>ancer genes) that can identify genetic and non-genetic drivers even with a limited number of samples. Applying APSiC to the large-scale deep shRNA screen Project DRIVE, APSiC identified well-known, pan-cancer genetic drivers, novel putative genetic drivers known to be dysregulated in specific cancer types and the context dependency of mRNA-splicing between cancer types. Additionally, APSiC discovered a median of 28 and 35 putative non-genetic oncogenes and tumor suppressor genes, respectively, for individual cancer types, including genes involved in genome stability maintenance and cell cycle. We functionally demonstrated that *LRRC4B*, a putative non-genetic tumor suppressor gene that has not previously been associated with carcinogenesis, suppresses proliferation by delaying cell cycle and modulates apoptosis in breast cancer.

**Conclusion:** We demonstrate APSiC is a robust statistical framework for discovery of novel cancer genes through analysis of large-scale perturbation screens. The analysis of DRIVE using APSiC is provided as a web portal and represents a valuable resource for the discovery of novel cancer genes.

## Background

Advances in large-scale functional screening technologies have enabled the discovery of gene requirements across diverse cancer entities [1,2]. Systematic perturbation screens using short hairpin RNA (shRNA) or CRISPR are increasingly used to investigate how genetic alterations or expression modulation of individual genes lead to phenotypic changes, revealing novel factors in carcinogenesis [3–5]. In parallel, large-scale sequencing efforts of 10,000+ cancers have provided a comprehensive molecular portrait of human cancers and their molecular pathogenesis [6]. Among the major findings is the unbiased discovery of genes mutated at rates significantly higher than the expected background level [7], revealing the global landscape of genetic ‘driver genes’ [8]. The discovery of these ‘driver genes’ forms the critical foundations of cancer diagnostics, therapeutics, clinical trial design and selection of rational combination therapies. Despite the large cohort size, the functional consequences of mutations in rarely mutated genes, such as *YAP/TAZ* [9], are only revealed by functional studies. Moreover, a systematic survey of non-genetic driver genes (i.e. driver genes for which the basis for oncogenicity is non-genetic) is lacking.

In a recent large-scale perturbation screen, the project DRIVE (deep RNAi interrogation of viability effects in cancer), 7,837 genes in 398 cancer cell lines were targeted with a median of 20 shRNAs per gene across a variety of malignancies to generate a comprehensive atlas of cancer dependencies [4]. The findings described in the study provided an overarching view of the nature and types of cancer dependencies, but has barely scratched the surface of the potential of the data generated. This huge resource of deep, robust and well-curated functional data, in conjunction of molecular profiling from the Cancer Cell Line Encyclopedia (CCLE) [10], remain largely untapped resources to be thoroughly mined and interrogated.

Analysis of shRNA perturbation screens is challenging due to the off-target effects of shRNAs as well as low number of cell lines screened [4,11]. Project DRIVE used deep coverage libraries to alleviate the off-target issues [4]. Additionally, several computational tools have been developed to elicit the on-target effect from a pool of siRNAs that have both on-and off-target effects [11–15]. RSA provides an absolute gene score based on the rank distribution of phenotypes produced by all shRNA reagents of a given gene [14]. ATARiS estimates a consensus shRNA profile for each gene and provides a relative gene score [13]. DEMETER is a regularized linear model for computing the effects due to knockdown of a target gene by taking into account the effects due to the seed sequence [11], while DEMETER2 is a complex hierarchical model structure that provides absolute gene dependency score as opposed to ATARiS and DEMETER that provide relative dependency score [15].

Cancer dependencies can be divided into two main groups of non-self dependencies (or synthetic lethality) and self-dependencies. Synthetic lethality refers to the cell dependency on the concomitant loss of two or more genes. In the context of an shRNA or CRISPR screen, synthetic lethality refers to the loss of cell viability upon knockdown/knockout of a gene conditional on the loss-of-function state of another gene. Various computational approaches have been used to discover synthetic lethal pairs from perturbation studies [16–18]. MiSL is a computational pipeline for the identification of synthetic lethal pairs using Boolean implications applied to the The Cancer Genome Atlas (TCGA) datasets [19,20]. A recent computational tool, called SLIdR, uses Irwin-Hall tests and causal inference for the discovery of pan-cancer as well as cancer-specific synthetic lethal pairs using the DRIVE data [21]. On the other hand, self-dependency refers to cell dependency on a gene with specific molecular features such as mutation, copy number, expression, and DNA methylation. Based on the molecular features, various statistical tests such as Fisher’s Exact and Wilcoxon tests, and differential expression analysis have been used to discover association between molecular features and gene dependencies. Most existing methods for finding self-dependencies are limited to the pan-cancer setting where a sufficient number of cell lines is available [4]. These tests lack statistical power for a limited number of observations (i.e. 5-10 observations) particularly after multiple testing corrections for a large number of tests. While the project DRIVE is one of the largest of its kind, the number of cell lines for some cancer types is quite small. A tool that can identify self-dependencies with small sample size is needed to make the best use of perturbation screens to reveal the cancer type-specific vulnerabilities.

Here, we introduce APSiC (<u>***A***</u>nalysis of <u>***P***</u>erturbation <u>***S***</u>creens for <u>***i***</u>dentifying novel <u>***C***</u>ancer genes), a novel tool for the systematic and robust interrogation of large-scale perturbation screens to discover gene (self-)dependencies for individual cancers even with limited number of samples. Incorporating mutation and copy number status of the samples, APSiC identifies potential genetic and non-genetic cancer genes. We consider three classes of genetic drivers, namely mutation oncogenes, amplification oncogenes, and mutation tumor suppressor genes as well as two classes of non-genetic drivers namely non-genetic oncogenes and tumor suppressor genes. Of particular importance, non-genetic drivers identified by APSiC have been largely less studied in cancer research. We applied APSiC to 26 cancers and identified both known and novel candidate genetic and non-genetic drivers. As a proof of concept, we functionally validated *LRRC4B* as a putative tumor suppressor gene in breast cancer. We provided the statistical analysis of DRIVE by APSiC as a web portal (https://apsic.shinyapps.io/APSiC/) for the scientific community to explore and functionally characterize genes that may be involved in carcinogenic processes and may pave the way for the discovery of novel cancer-related biomarkers and drug targets.

## Results and Discussion

### The APSiC Algorithm

APSiC uses rank-based statistics to discover self-dependencies in perturbation screens (see Methods for a technical description of the algorithm). Given the raw cell viability readout of a perturbation screen, APSiC computes a rank profile for each gene by first ranking all genes by their viabilities upon knockdown in a given sample to the range of [0, 1] then aggregating the normalized ranks for a given gene across all samples (Fig. 1a). Thus ranks close to zero represent reduced viability while the ranks close to one indicate cell growth upon knockdown. Incorporating mutation and copy number status of the samples, APSiC identifies potential genetic and non-genetic cancer genes by assessing deviation of the distribution of normalized ranks from what is expected by chance using the Bates and Irwin-Hall tests. The Irwin-Hall distribution has been successfully used in the identification of synthetic lethal gene pairs [21] and prioritization of cancer genes based on multi-omics data [22]. The use of the rank-based statistics with the Bates and Irwin-Hall distributions provides enhanced statistical power when the number of cell lines is limited.

**Fig. 1.**
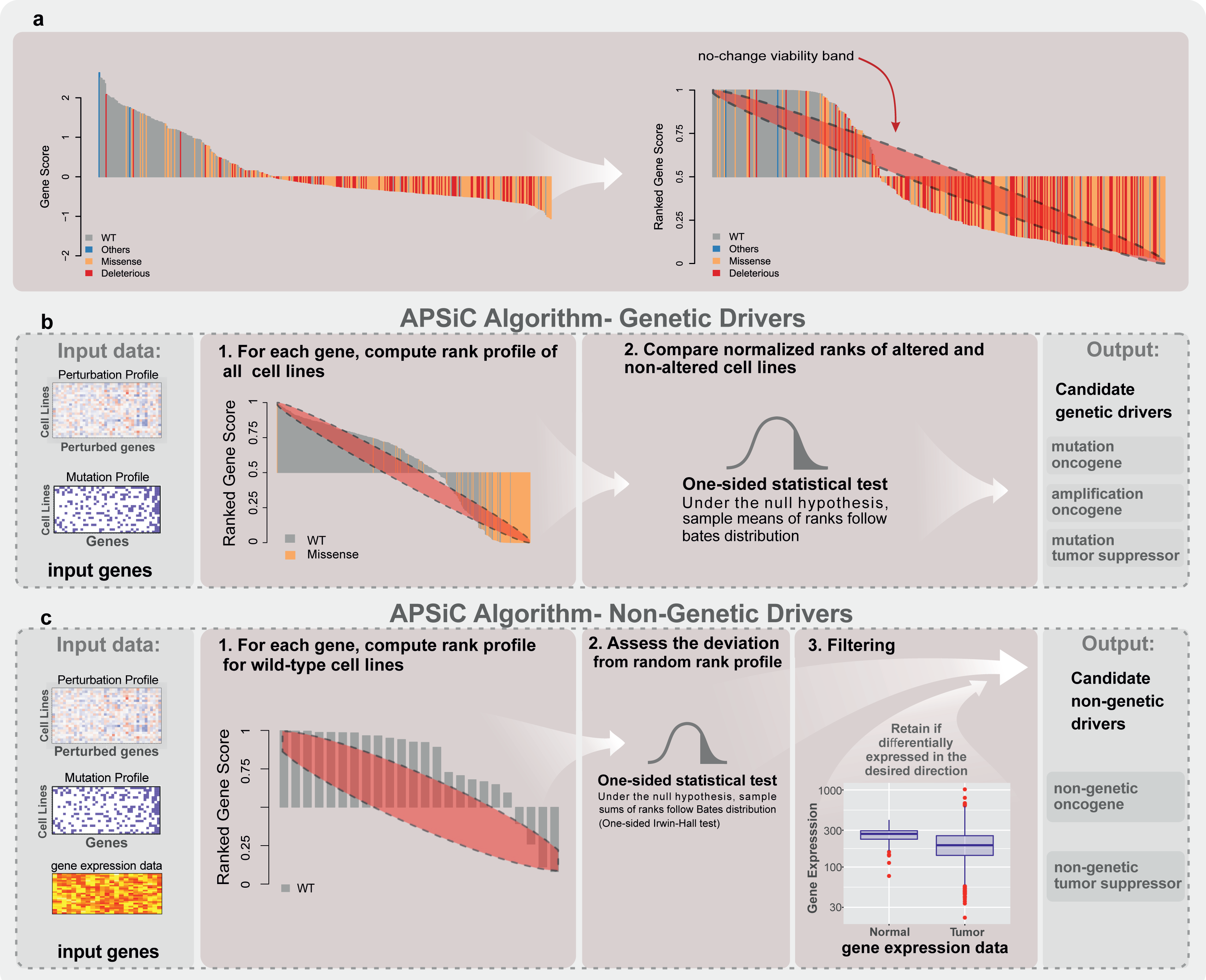
Overview of the APSiC algorithm. **a**, Illustration of the transformation from the raw cell viability scores for a given gene (left, using *TP53* as an example) to the rank profile (right). Each bar of the waterfall plots represents one sample and is colored by the mutational status of the given gene in the sample. The red ellipse in the rank profile (right) represents a no-change (random) viability band. **b-c**, Schematic representation of the APSiC algorithm for identifying **(b)** genetic and **(c)** non-genetic drivers (See Materials and Methods for details).

We consider three classes of genetic drivers (mutation oncogenes, amplification oncogenes, and mutation tumor suppressor genes) and two classes of non-genetic drivers (non-genetic oncogenes and tumor suppressor genes). We define mutation and amplification oncogenes as genes for which reduced cell viabilities are preferentially observed in samples with missense mutations and copy number amplifications, respectively, while mutation tumor suppressor genes are those for which increased viabilities are preferentially observed in samples with deleterious mutations. To identify such genetic drivers, we test, for a given gene, whether ranks of the samples with and without the specific class of genetic alteration are significantly different using a one-sided Bates test (Fig. 1b, **Additional files 1-3**). For mutation and amplification oncogenes, we compute the lower-tailed *P* values (i.e. the ranks preferentially suggest reduced viability upon gene knockdown), while for mutation tumor suppressor genes, we compute the upper-tailed *P* values (i.e. the ranks preferentially suggest increased cell viability upon knockdown). For the non-genetic drivers, we test whether gene knockdown in samples without genetic alteration in the gene has any impact on cell viability by computing lower and upper-tailed Irwin-Hall test *P* values for oncogenes and tumor suppressor genes, respectively (Fig. 1c, **Additional files 4-5**). Optionally, we further test whether the expression of candidate non-genetic oncogenes or tumor suppressor genes is respectively enhanced or repressed in human tumors compared to the corresponding normal tissue type.

### APSiC identifies well-known and novel genetic cancer drivers

We applied APSiC to the DRIVE perturbation screens and the genetic data from the CCLE [10] to identify genetic driver genes (**Additional files 1-3**). The dataset consists of 383 cell lines across 26 cancer types, with a median of 11 (range 5-40) cell lines per cancer type (Fig. 2a). In a pan-cancer analysis, APSiC reassuringly identified the well-known mutation oncogenes *BRAF, CTNNB1, KRAS, NRAS, PIK3CA* and *TP53* (Figs. 2b-c, **Additional file 6**). Additionally, *DDX27, DCAF8L2* and *RBM39* were detected as mutation oncogenes (**Additional files 6 and 7**). The top amplification oncogene were *KRAS, BRAF, CDK4, YAP1, IL6* and *HAS2* (Figs. 2b, d and **Additional file 6**), while the only mutation tumor suppressor was *ARID1A* (Figs. 2b, e and **Additional file 6**). However, the identification of mutation tumor suppressor genes in a knockdown screen is likely to have limited utility given that mutation tumor suppressor genes are frequently associated with loss of the wild-type allele.

**Fig. 2.**
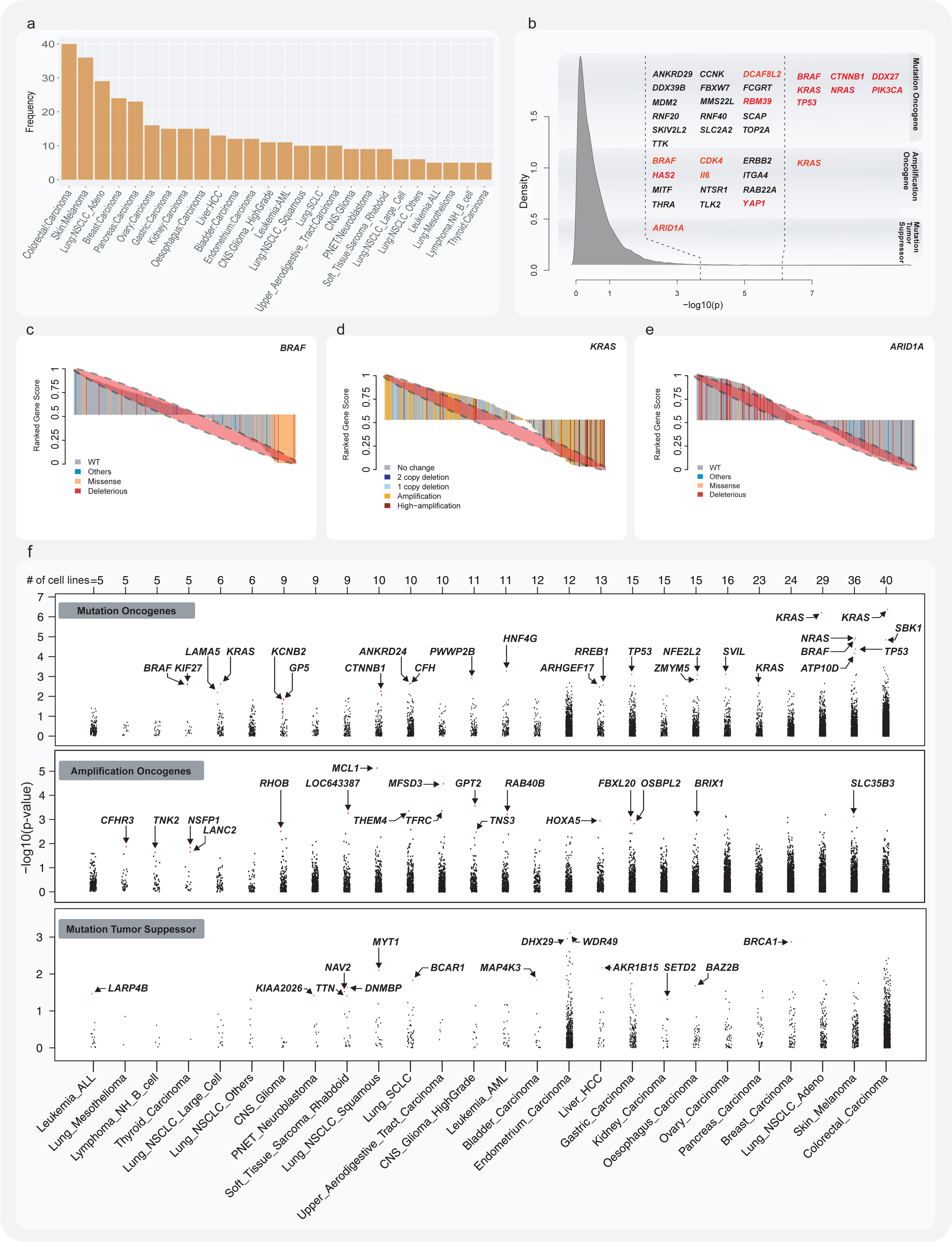
Analysis of the genetic drivers in the DRIVE perturbation screen. **a**, The number of cell lines available for 26 cancer subtypes in the DRIVE perturbation screen. **b**, Kernel density estimation of the *P* values (on a -log_10_ scale) for genetic drivers using the APSiC algorithm in a pan-cancer analysis. Candidate mutation oncogenes, amplification oncogenes, and mutation tumor suppressors identified by the APSiC are shown, with genes reaching significance level after multiple testing corrections highlighted in red. **c**, Rank profile for a mutation oncogene (*BRAF*) colored by mutation status. **d**, Rank profile of an amplification oncogene (*KRAS*) colored by copy number status. **e**, Rank profile of a mutation tumor suppressor (*ARID1A*) colored by mutation status. **f**, Dot plots of the *P* values (on a -log_10_ scale) for genetic drivers using the APSiC algorithm in a cancer type-specific analysis, for (top) mutation oncogenes, (middle) amplification oncogenes and (bottom) mutation tumor suppressors. Genes reaching significance level after multiple testing corrections are highlighted in red. Cancer types are sorted by the number of cell lines.

One of the main strengths of APSiC is the identification of dependencies in small sample sets. We therefore applied APSiC to the DRIVE data to identify genetic driver genes for individual cancer types. Across the 26 cancer types, we found 20, 18 and 14 mutation oncogenes, amplification oncogenes and mutation tumor suppressor genes, respectively (Fig. 2f and **Additional files 1-3**), in 15, 14 and 11 cancer types. We found no correlation between the number of cell lines and the number of driver genes identified (p=0.14, 0.21 and 0.97 for mutation oncogenes, amplification oncogenes and mutation tumor suppressor genes, respectively, Spearman correlation tests). *KRAS*, *BRAF* and *TP53* were identified as a mutation oncogene in 4, 2 and 2 cancer types, respectively, while the other 17 genes were identified as mutation oncogenes in a single cancer type each. Other well described mutation oncogenes include *NFE2L2* in esophagus carcinoma and *CTNNB1* in non-small cell lung cancer. The remaining putative mutation oncogenes have not been reported as frequently mutated in human cancers. These putative novel mutation oncogenes include *RREB1* (Ras Responsive Element Binding Protein 1) and *SBK1* (SH3 Domain Binding Kinase 1), both of which have previously been found to be dysregulated in cancer [23,24].

No amplification oncogene or mutation tumor suppressor was identified in more than one cancer type. We identified *MCL1* and *RHOB* as the top amplification oncogenes in the squamous subtype of non-small cell lung cancer and glioma, respectively (Fig. 2f and **Additional files 2-3**). *HOXA5*, a gene frequently overexpressed in hepatocellular carcinoma [25], was identified as an amplification oncogene. Of the mutation tumor suppressor genes, *BRCA1* and *SETD2* were found in breast and kidney cancers, respectively. *BAZ2B*, a gene involved in chromatin remodeling, is a putative mutation tumor suppressor in esophageal cancer. Of note, 15% of *BAZ2B* somatic mutations in the TCGA pan-cancer cohort are truncating mutations [8], suggestive of a tumor suppressor role for *BAZ2B.* Taken together, APSiC identified both well known and putative genetic driver genes previously linked to carcinogenesis.

### Survey of non-genetic driver genes reveals cancer type specificity

By identifying strong irregular viability patterns in the rank profiles of cell lines wild-type for a given gene using APSiC, we evaluated the non-genetic dependencies across cancer types in the DRIVE data (**Additional files 4-5**). Consensus clustering of the most variable genes in terms of APSiC *P* values revealed that such non-genetic dependencies segregate by organ systems or cell-of-origin into four clusters (Fig. 3a). In particular, non-epithelial cancers including leukemias/lymphomas, sarcomas, gliomas and neuroblastomas form a cluster distinct from epithelial cancers including those of the lungs, the breasts and gastrointestinal tract. This is consistent with the observation that multi-omics cancer classification is primarily driven by cell-of-origin and anatomic regions [26]. Furthermore, the top-level segregation of the cancer types was largely driven by the context-dependency of mRNA-splicing genes. We observed that mRNA-splicing genes such as *PRPF6* (Pre-MRNA Processing Factor 6) and *SART3* (Spliceosome Associated Factor 3, U4/U6 Recycling Protein) were tumor suppressive in the cluster enriched for non-epithelial cancers while they were oncogenic in the epithelial cancer cluster (Fig. 3b). The context-dependency highlights the divergent role of mRNA-splicing in carcinogenesis between cancer types. Our results also underscore the necessity for an algorithm powerful enough to analyze perturbation screens for small numbers of samples in a cancer type-specific manner.

**Fig. 3:**
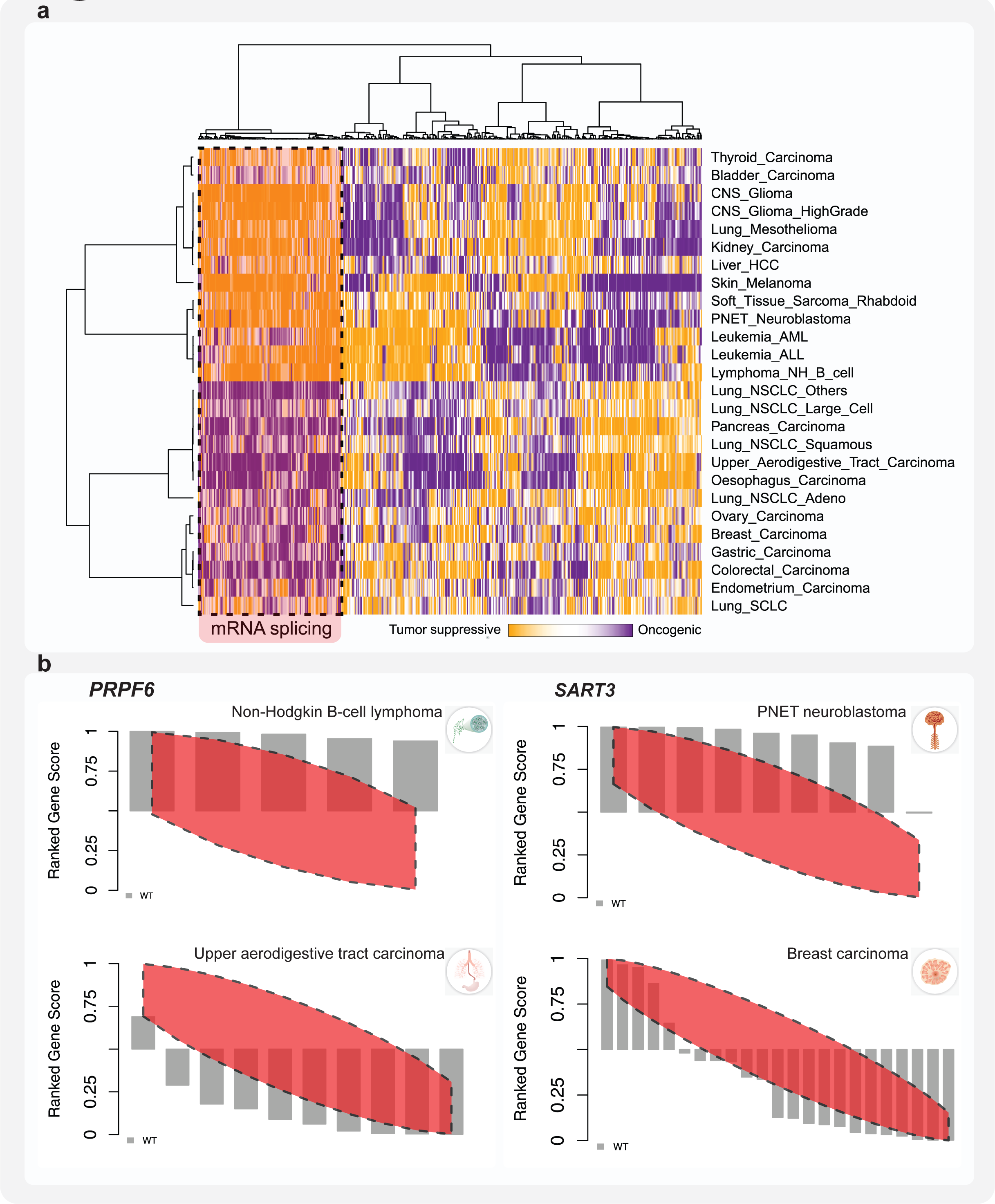
Pan-cancer analysis of the non-genetic drivers in the DRIVE perturbation screen. **a**, Heatmap illustrates consensus clustering of the 500 most variable *P* values for non-genetic drivers using the APSiC algorithm for the 26 cancer types in the DRIVE perturbation screen. Consensus clustering identified 4 clusters across cancer types and 5 clusters across the 500 genes. One of the clusters was enriched for genes involved in mRNA splicing and processing. **b**, Rank profiles for *PRPF6* in (top) non-Hodgkin B-cell lymphoma and (bottom) upper aerodigestive tract carcinoma and for *SART3* in (top) PNET neuroblastoma and (bottom) breast carcinoma.

Based on the DRIVE screen alone, we identified a median of 28 non-genetic oncogenes (range 6-557) and 35 non-genetic tumor suppressor genes (range 1-471) per cancer type. However, we reasoned that the many non-genetic onco- and tumor suppressor genes would also be over- and under-expressed, respectively, in the corresponding cancer types. For the 12 cancer types for which gene expression data for the cancer and corresponding non-cancer counterparts were available from the TCGA (**Additional file 8**), we further restricted the putative non-genetic onco- and tumor suppressor genes to those that were over- and under-expressed, respectively, relative to their non-cancer counterparts. After this filtering step, there were a median of 13 non-genetic oncogenes (range 2-117) and 3 non-genetic tumor suppressor genes (range 0-42, Fig. 4) per cancer type. We identified several well-known oncogenes, including *CDK1* (a master regulator of cell cycle) and *SMC1A* (a component of the cohesin complex involved in cell cycle checkpoint and genome stability)[27], and some that have been shown to have oncogenic properties in some cancer types, such as *MKI67IP* (or *NFIK*) [28]. We also identified *TEAD3*, a lesser described member of the TEAD family involved in hippo signalling, as oncogenic in liver cancer [29]. Among the top candidate tumor suppressors were *FOXP2* in endometrial cancer and *XRCC5* in kidney carcinoma. *FOXP2* knockdown has been shown to promote tumor initiation and metastasis in breast cancer [30] while *XRCC5*, encoding the protein Ku80, is a key DNA damage repair protein. However, we also identified many genes that have not been associated with carcinogenesis.

**Fig. 4:**
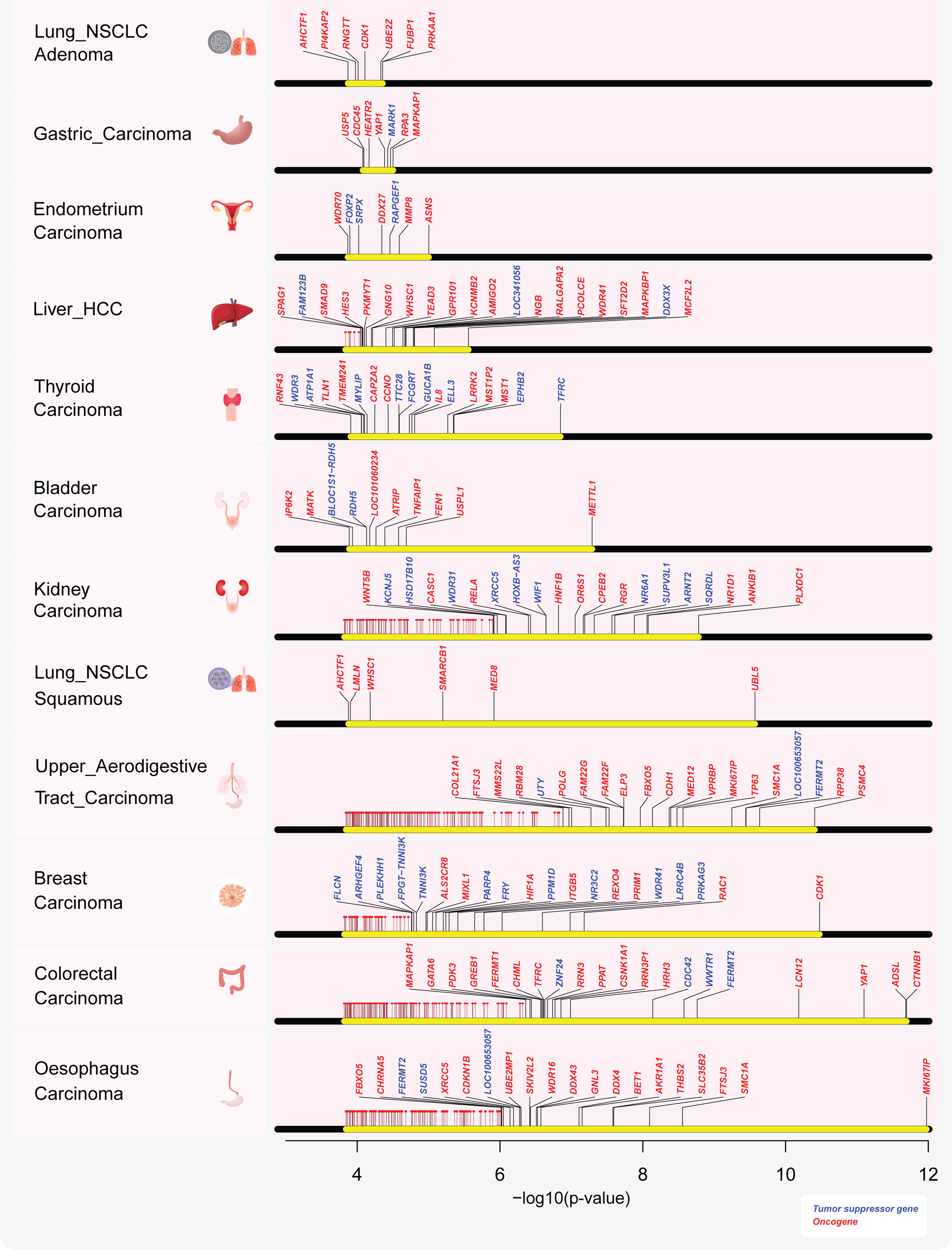
Cancer type-specific analysis of the non-genetic drivers in the DRIVE perturbation screen. Non-genetic driver genes identified in the 12 cancer types with corresponding gene expression data from the TCGA. APSiC P values are shown (on a -log_10_ scale) for driver genes significant after multiple testing corrections and over- or underexpressed in human cancers for oncogenes (in red) and tumor suppressor genes (in blue), respectively. The top 20 genes for each cancer type are labelled.

### *LRRC4B* is a putative tumor suppressor gene in breast cancer

As a proof-of-concept to validate APSiC, we selected *LRRC4B*, the top putative non-genetic tumor suppressor gene in breast cancer, for functional validation. A literature search of *LRRC4B* in cancer suggests that its role and function in carcinogenesis are unknown. Meanwhile, one of its paralogs *LRRC4* has been shown to have a putative tumor suppressor role in glioma by modulating the extracellular signal-regulated kinase/protein kinase B/nuclear factor-κB pathway [31]. In the DRIVE RNAi screen, nearly all breast cancer cell lines displayed significantly increased cell viability upon *LRRC4B* knockdown and breast cancers in TCGA showed lower *LRRC4B* expression compared to normal breast tissue (**Additional file 9a**). We selected the breast cancer cell lines MDA-MB231, BT-549 and MCF-7 with high, moderate and low endogenous *LRRC4B* expression to investigate whether *LRRC4B* knock-down would result in other classical cancer phenotypes such as increased migration and colony formation (**Additional file 9b**). We silenced *LRRC4B* in MDA-MB231 and BT-549 using siRNA, reducing LRRC4B protein expression by 40% and 60%, respectively, 72 hours post-transfection (Figs. 5a, e). In both models, *LRRC4B* downregulation significantly increased the proliferation and migration rates (Figs. 5b-c, f-g). By contrast, *LRRC4B* overexpression significantly reduced proliferation and migration in MCF-7 (Figs. 5i-k).

**Fig. 5:**
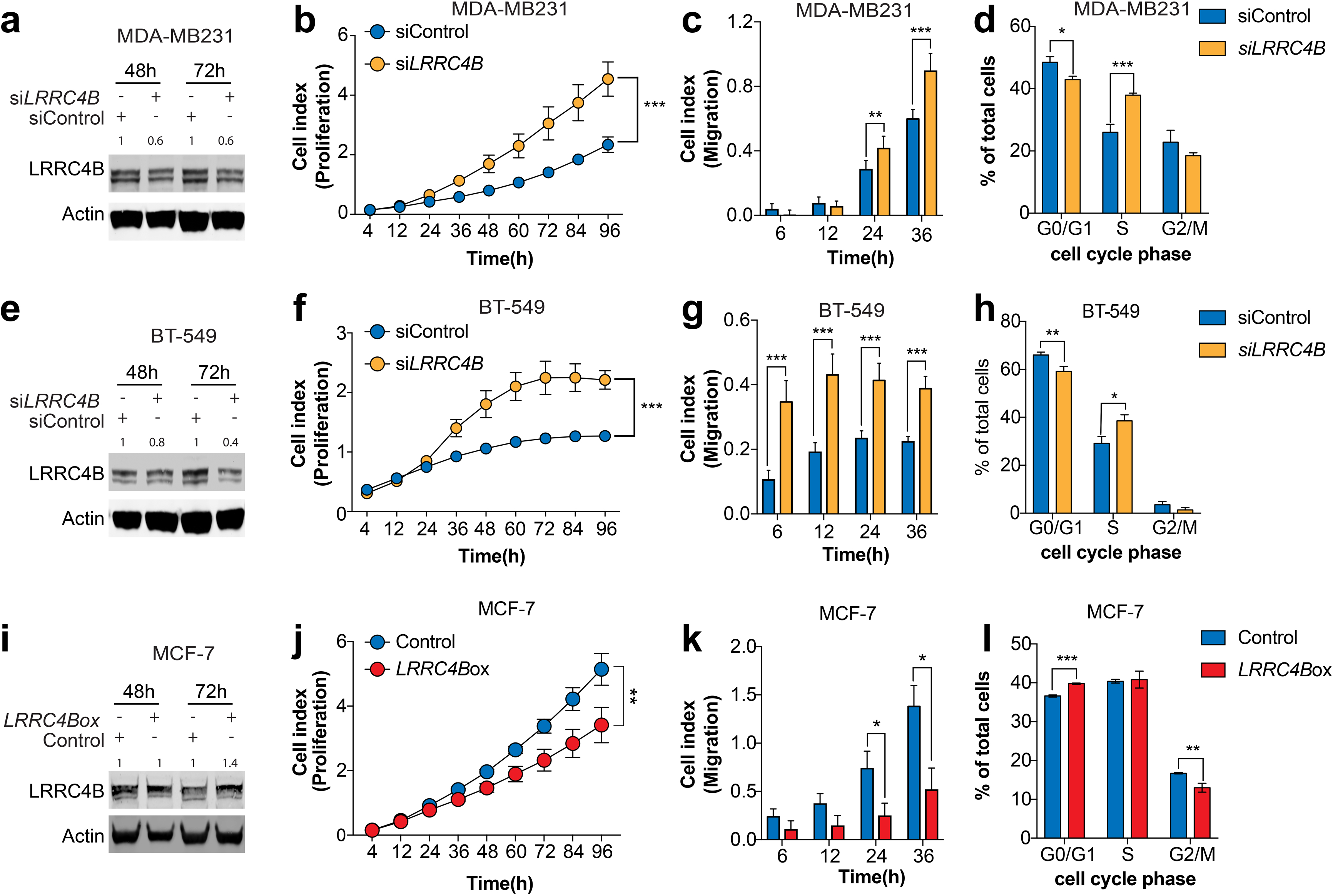
*LRRC4B* has tumor suppressor-like properties in *in-vitro* models of breast cancer. **a, e, i**, Western blotting showing LRRC4B protein level in (**a**) MDA-MB231, (**e**) BT-549 and (**i**) MCF-7 cell lines 48 and 72 hours post transfection. Actin was used as a loading control and for normalization. **b, f, j**, Proliferation kinetics of (**b**) MDA-MB231, (**f**) BT-549 and (**j**) MCF-7 cells upon (**b, f**) downregulation or (**j**) upregulation of *LRRC4B* compared with the control. **c, g, k**, Migration potential of **(c)** MDA-MB231, (**g**) BT-549 and (**k**) MCF-7 cells upon (**c, g**) downregulation or (**k**) upregulation of *LRRC4B* compared with the control. **d, h, i**, Cell cycle analysis of (**d**) MDA-MB231, (**h**) BT-549 and (**i**) MCF-7 cells upon (**d, h**) downregulation or (**i**) upregulation of *LRRC4B* compared with the control. Error bars represent standard deviation obtained from three independent experiments. For all experiments, statistical significance was assessed by multiple t-tests (* *P* < 0.05, ** *P* < 0.01, *** *P* < 0.001).

*LRRC4* has been shown to suppress cell proliferation by delaying cell cycle in late G_1_ phase [32],[33]. To test whether *LRRC4B* may play the same role in breast, we analyzed cells with *LRRC4B* overexpression or downregulation stained with DAPI by flow cytometry (FACS). *LRRC4B* knockdown in MDA-MB231 and BT-549 promoted cell transition into S phase (Figs. 5d, h), while *LRRC4B* overexpression in MCF-7 significantly retained cells in G_1_ phase (Fig. 5l), suggesting a similar mechanism to *LRRC4*.

A common mechanism of oncogenicity is resistance to apoptosis [34]. To test whether modulation of apoptosis is a mechanism of action of *LRRC4B* as an oncosuppressor, we induced apoptosis with doxorubicin and measured it using Annexin V and propidium iodide co-staining followed by FACS analysis (Fig. 6a). Forty eight hours after treatment, *LRRC4B*-overexpressing MCF-7 cells showed 10% more apoptotic and 10% fewer live cells, suggesting that *LRRC4B* overexpression could sensitize cells to doxorubicin-induced apoptosis (Fig. 6b). By contrast, *LRRC4B-*downregulating MDA-MB231 and BT-549 cells showed increased resistance to doxorubicin and had 25% and 10% fewer apoptotic and 25% and 10% more live cells, respectively (Fig. 6b). Our results provide compelling evidence that APSiC identified *LRRC4B* as a novel oncosuppressor gene in breast cancer.

**Fig. 6:**
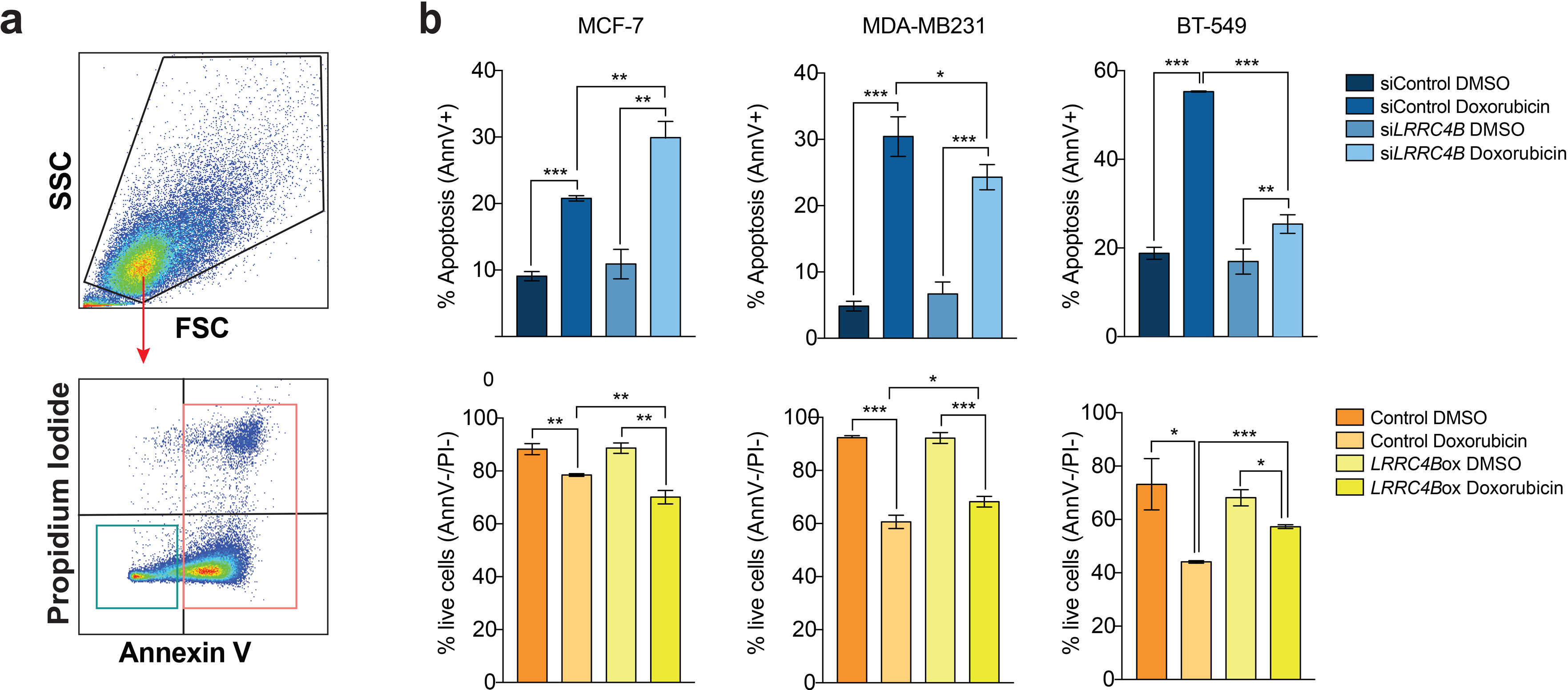
*LRRC4B* expression modulates induction of apoptosis by Doxorubicin. **a**, Dot plot illustrating the flow cytometry gating strategy used to assess cell viability and apoptosis using Annexin V and propidium iodide staining of MDA-MB231, BT-549 and MCF-7 cells upon downregulation/ upregulation of *LRRC4B* compared with the control cells, with and without Doxorubicin. **b**, Quantification of the mean (+/-SD) percentage of apoptotic cells (AnnV+) and live cells (AnnV-/PI-) across the different groups. Error bars represent standard deviation obtained from three independent experiments. For all experiments, statistical significance was assessed by multiple t-tests (* *P* < 0.05, ** *P* < 0.01, *** *P* < 0.001).

## Conclusions

APSiC is a novel tool to enable systematic and robust discovery of gene dependencies from small number of samples and can conceivably be applied to other types of perturbation screens, such as CRISPR screens. Our analysis of the DRIVE perturbation screens using APSiC is a valuable resource for the discovery of drug targets, cancer-related biomarkers and novel cancer genes, in particular of non-genetic driver genes for which a systematic analysis has been lacking. Our results would complement the associated multi-omics profiling [35,36] and drug screens [37] to understand the vulnerabilities of cancer.

## Methods

### The detailed explanation of the APSiC algorithm

In this section, we give a more technical description of APSiC and also introduce a new waterfall plot, called rank profile, for visualization of gene dependencies. First we briefly describe some necessary definitions and background material from ordered statistics. We consider the knockdown experiments of *p* genes across *N* cell lines. Let *ν*_*ij*_ be viability of cell line *i∈{1,…,N}* upon knocking down gene *j∈{1,…, p}* and *m*_*ij*_ be a binary variable indicating whether a specific genetic alteration (i.e. mutation or copy number alteration) is present in gene *j* of cell line *i*. In this study, we only consider deleterious (e.g. nonsense, frameshift, splice site and mutations affecting start or stop codons) and missense mutations. Waterfall plots are often used to show viabilities of knockdown experiments for a single gene across different cell lines and are aimed to illustrate different gene dependencies. As an example, waterfall plot for gene *TP53* is shown in Fig. 1a (left). Each vertical bar corresponds to a cell line and is colored by the pre-existing mutation types present in *TP53*. Fig. 1a indicates cell lines with the presence of deleterious or missense mutations in *TP53* tend to have lower viabilities upon knockdown of this gene. While waterfall plot is a useful visualization tool for demonstrating gene dependencies, it lacks sufficient interpretability in certain cases, particularly when the number of cell lines is limited. In this paper, we introduce a new waterfall plot, named rank viability profile or simply rank profile, to address this issue.

To make viability scores comparable across cell lines, we compute normalized rank values per cell lines denoted as, representing the rank of viability for gene *j* among all knockdown experiments in cell line *i*. For mathematical convenience and without loss of generality, we normalized ranks to the range of [0, 1]. When the number of knockdown genes is high, normalized ranks have many distinct levels in the interval [0, 1] and we assume normalized ranks are continuous. Let denote random variables associated to ranks of a gene *A* in *N* cell lines. We drop subscript A and denote ranks as for the simplicity of notation. By placing ranks,, in ascending orders and renaming them, we obtain where is called *i*th ordered statistic. It is easy to see that and. The probability density function of ordered statistic in general is given as

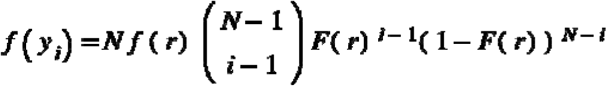

where *f(r)* and *F(r)* denote probability density and cumulative distribution functions, respectively. If there is no dependency between knocking down of a gene and the viability of the cell, we can assume for, hence we have. Using this result, we can construct a no-change viability band at statistical significance α using the quantiles of at the and for. Now we define a new waterfall plot, called rank viability profile or simply rank profile, as a waterfall plot using normalized ranks, realizations of for a gene, overlaid with no-change viability band (Fig. 1a).

The APSiC algorithm identifies potential cancer genes by assessing deviation of respective rank profiles from what is expected by chance. The algorithm can identify both genetic and non-genetic drivers (Fig. 1b-c). We consider three categories for genetic drivers.

- Mutation oncogene: defined as genes for which reduced viabilities are observed preferentially in samples with missense mutation.
- Amplification oncogene: defined as genes for which reduced viabilities are observed preferentially in samples with copy number amplification.
- Mutation tumor suppressor: defined as genes for which increased viabilities are observed preferentially in samples with deleterious mutation.

We consider two categories for non-genetic drivers, namely

- Non-genetic oncogene: defined as genes for which reduced viabilities are observed in samples without a genetic alteration in the respective gene.
- Non-genetic tumor suppressor: defined as genes for which increased viabilities are observed preferentially in samples without a genetic alteration in the respective gene.

For genetic drivers, the APSiC algorithm considers rank profiles of mutated and wild-type samples with respect to an input gene *g* (Fig. 1b). Then, it performs a one-sided statistical test to determine whether rank scores of the two groups of samples are significantly different in the direction of interest, according to the genetic feature of interest. Suppose 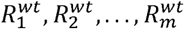 and 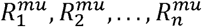 are random variables denoting rank scores upon knockdown of gene *g* for *m* wild-type and n mutated samples, respectively. Let 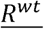 and 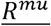 denote the average of ranks for the wild-type and mutated samples, respectively. We define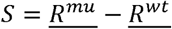 as the test statistic and *s*_*obs*_ as the observed test statistic. We assume the null hypothesis is that the knockdown of gene *g* does not have any impact on the viability of samples and therefore there is no difference in average of ranks for two groups, i.e. *S* =0. The general formula for the distribution of any weighted sum of uniform random variables is given in Kamgar-Parsi, 1994[38]. We simplify the general formula thereby and obtain the null distribution of the test statistic *S*as

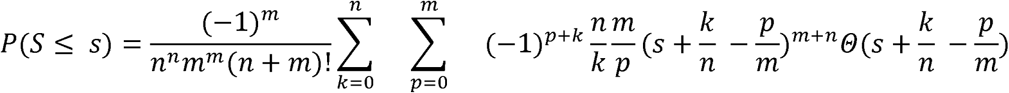

where

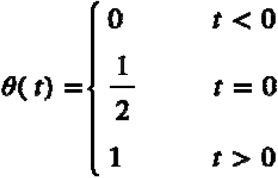

For mutation and amplification oncogenes, we compute lower-tailed *P* values, while for mutation tumor suppressors, we compute upper-tailed *P* values,. Due to numerical issues, it is impractical to use the exact null distribution formula for large values of *m* and *n* (*m+n* >20). In this case, we compute an approximation for the null distribution of *S* as follows. Under the null hypothesis, —— and —— follow *Bates* distributions *Bates(m)* and *Bates(n)*, respectively. The Bates distribution is a distribution that represents the mean of a number of independent uniform random variables on the unit interval. For large values of *m* and *n*, —— and —— are approximately distributed – —— by and – ——. Hence, under the null hypothesis the test statistic *S* is approximately distributed as — — –.

To identify non-genetic drivers, we only consider the wild-type (i.e. without non-synonymous mutations and without copy number amplification (GISTIC copy number state 2) or deep deletions (GISTIC copy number state −2)) samples with respect to an input gene (Fig. 1c). The null hypothesis is that the knockdown of a gene *g* does not have any impact on the viability of the samples. We define the test statistic as and as the observed test statistic. Under the null hypothesis, *T* follows an Irwin-Hall distribution, which represents the summation of independent uniform random variables on the unit interval. For large values of *m*, *S* is approximately distributed as To identify non-genetic oncogenes, we require significant lower-tailed *P* values, for wild-type cell lines with respect to the input gene. Additionally, for the respective tissue type, the overall expression at the RNA level of a putative oncogene in tumor samples is required to be significantly higher than the one in normal tissue samples using the *t* test. On the contrary, for identifying non-genetic tumor suppressors, we require significant upper-tailed *P* values, *P*(*T ≥ t*_*obs*_), for wild-type cell lines with respect to the input gene as well as lower RNA expression of tumor samples in comparison to normal tissue samples using the *t* test.

### Downloading and preprocessing of DRIVE and TCGA data

We considered the viability profiles of 383 cell lines in the project DRIVE [4] for which their genetic profiles were available at the Cancer Cell Line Encyclopedia (Fig. 2a) [10]. We computed aggregated gene-level viability scores for each experiment by the RSA and ATARiS algorithms [13,14]. The RSA and ATARiS scores are available for 7726 and 6557 genes, respectively. We used the same method as defined in the project DRIVE to remove essential genes, defined as genes with an RSA value of ≤ 3 in more than half of cell lines [4]. After the removal of 184 essential genes by this method, for pan-cancer analysis, the ATARiS scores are available for 6373 genes in 383 cell lines in 39 cancer types over 17 primary tissues. For the genetic drivers, we considered only genes for which there are at least 2 samples harbored a genetic alteration of the corresponding class in individual cancer types. For the non-genetic drivers, we only considered genes for which there are at least 2 samples wild-type for the gene in individual cancer types. For pan-cancer analysis, the above threshold was at least 4 samples for both genetic and non-genetic drivers. For the identification of mutation tumor suppressor genes, mutations annotated as *In_Frame_Ins*, *In_Frame_Del*, *Frame_Shift_Ins*, *Frame_Shift_Del*, *Nonsense_Mutation*, *Splice_Site*, *Start_Codon_Del*, *Stop_Codon_Del*, *Stop_Codon_Ins*, *Start_Codon_Ins* were considered deleterious. For the analysis of individual cancer types, we considered the 26 cancer types for which more than four cell lines are available in the DRIVE data.

TCGA gene expression data was obtained for 12 cancers for which data are available for tumor and normal tissues (**Additional file 8**). The data were downloaded using the TCGAbiolinks package in R [39]. The normalized expression level of genes in CPM (counts per million) were used for the identification of non-genetic drivers.

### Multiple testing

To address the multiple comparisons problem, we chose a significance level such that the expected number of false positives due to multiple testing for each cancer and feature is equal to one. To this end, we chose a significance level of 1/*n*, or 0.05 if 1/*n*>0.05, where *n* is the number of genes tested for identification of drivers. Using this approach, we were able to keep many interesting hits while keeping the number of false positive cases low.

### Clustering and pathway analysis

Clustering was performed using ConsensusClusterPlus Bioconductor package in R [40] using 1-Spearman correlation as the distance metric and the Ward hierarchical clustering algorithm. The number of clusters was determined based on the relative change in area under the consensus cumulative distribution function over the number of evaluated clusters. Pathway analysis was performed using g:Profiler [41].

### Cell lines

Breast cancer derived cell lines (MCF-7, BT-549 and MDA-MB231) were maintained in a 5% CO_2_-humidified atmosphere at 37°C and cultured in DMEM supplemented with 10% FBS, 1% Pen/Strep (Bio-Concept) and 1% MEM-NEAA (MEM non-essential amino acids, ThermoFisher Scientific). All cell lines were confirmed negative for mycoplasma infection using the PCR-based Universal Mycoplasma Detection kit (American Type Culture Collection, Manassas, VA) as previously described [42].

### Transient gene knockdown by siRNAs

Transient gene knockdown was conducted using ON-TARGET plus siRNA transfection. ON-TARGET plus SMARTpool siRNAs against human *LRRC4B* (Dharmacon, CO; #L-023786-01-0005), ON-TARGET plus SMARTpool non-targeting control and DharmaFECT transfection reagent (Dharmacon, CO; #T-2001-03) were all purchased from GE Dharmacon. Transfection was performed according to the manufacturer’s protocol. Briefly, log-phase breast cancer cells were seeded at approximately 60% confluence. Because antibiotics affects the knockdown efficiency of ON-TARGET plus siRNAs, growth medium was removed as much as possible and replaced by antibiotic-free complete medium. siRNAs were added to a final concentration of 25 nM. Cells were incubated at 37°C in 5% CO_2_ for 24-48-72 hours for 48-72 hours for protein analysis. To avoid cytotoxicity, transfection medium was replaced with complete medium after 8 hours.

### Protein extraction and western blot

Proteins were extracted using Co-IP buffer (100 mmol/L NaCl, 50 mmol/L Tris pH 7.5, 1 mmol/L EDTA, 0.1% Triton X-100) supplemented with 1x protease inhibitors (cOmplete Mini, EDTA-free Protease Inhibitor Cocktail, Roche, CO, #4693159001) and 1x phosphatase inhibitors (PhosSTOP #4906837001, Merck). Cell lysates were then treated with 10x reducing agent (NuPAGE Sample Reducing Agent, Invitrogen, #NP0009), 4x loading buffer (NuPAGE LDS Sample Buffer, Invitrogen, #NP0007), boiled and loaded into neutral pH, pre-cast, discontinuous SDS-PAGE mini-gel system (NuPAGE 10% Bis-Tris Protein Gels, ThermoFisher). The proteins were then transferred to nitrocellulose membranes using Trans-Blot Turbo Transfer System (Bio-Rad). The membranes were blocked for 1 hr with Sure Block (Lubio Science) and then probed with primary antibodies overnight at 4°C. Next day, the membranes were incubated for 1 hr at RT with fluorescent secondary goat anti-mouse (IRDye 680) or anti-rabbit (IRDye 800) antibodies (both from LI-COR Biosciences). Blots were scanned using the Odyssey Infrared Imaging System (LI-COR Biosciences) and band intensity was quantified using ImageJ software. The ratio of proteins of interest/loading control in treated samples were normalized to their counterparts in control cells. Antibodies against LRRC4B (PA5-23529, Thermofisher) and B-actin (A5441, Sigma) were used at dilution 1:1000 and 1:5000, respectively.

### Proliferation assay

Cell proliferation was assayed using the xCELLigence system (RTCA, ACEA Biosciences, San Diego, CA, USA) as previously described.[43] Cells were first seeded and transfected in 6 well plates and 24 h after transfection 5×10^3^ cells were resuspended in 100 μl of medium and plated in each well of an E-plate 16. Background impedance of the xCELLigence system was measured for 12 s using 50 μl of room temperature cell culture media in each well of E-plate 16. The final volume in each well was then 150 μl. The impedance signals were recorded every 15 minutes until 96/120 h and expressed as cell index values, calculated automatically and normalized by the RTCA Software Package v1.2. The values were defined as mean ± standard deviation. Mann-Whitney test was used for statistical analysis with GraphPad software.

### Migration assay

Migration assays were performed using the CIM-plate of the xCELLigence Real-Time Cell Analysis (RTCA, ACEA Biosciences, San Diego, CA, USA) system. Cells were first transfected in 6-well plates and 24 h after transfection, they were harvested and seeded in the CIM-plate. Every well of the bottom chamber was filled with 160 μl of the corresponding medium at 10% FBS concentration. After placing the upper chamber on top of the lower chamber, 50 μl of serum free medium was added on each CIM well for the background measurement. After 3x PBS washing, 3×10^4^ cells re-suspended in 100 μl of the corresponding medium at 1% FBS concentration were seeded in each well of the upper chamber. The measurements were taken every 15 minutes until 24 h after seeding and expressed as cell index values. Mann-Whitney U test was used for statistical analysis with GraphPad software.

### Cell cycle analysis

Seventy-two hour after transfection, cells were collected, stained with DAPI and analyzed by flow cytometry using the BD FACS Canto II cytometer (BD Biosciences, USA). Briefly cells were harvested and washed 2X in PBS to get rid of serum proteins at 1200 rpm for 5 minutes. Pellets (up to 3×10^6^ cells) were resuspended in 1.2 ml PBS (Ca and Mg free). For crosslinking proteins 3.0 ml of 95% ice cold EtOH was added dropwise while vortexing. Cells were fixed in this final 70% Et-OH solution for at least 30 minutes or over night. The Et-OH/cell suspension was then diluted with 12 ml of PBS (for a total volume of 15 ml) and centrifuge at 2000-2200 rpm for 10 min. Cells were then washed once more with 15 ml PBS and then resuspended in 0.5-2.0 ml of DAPI stain solution (0.1% TritonX 100 and 10 ug/ml). After 30 min of incubation on ice cells were analyzed by flow cytometry, measuring the fluorescence emission at 461 nm. Data were analysed using the FlowJo software version 10.5.3 (https://www.flowjo.com).

### Apoptosis analysis by flow cytometry

BT-549 and MDA-MB231 cells were transfected with siRNA (control or against *LRRC4B*) and MCF-7 cells were transfected with *LRRC4B* overexpressing plasmid or control plasmid. Eight hours after transfection medium was changed and doxorubicin added according to the respective IC50 for each cell line[44]^,[45]^. Cells were collected 60 hours post siRNA transfection or *LRRC4B* overexpression and 48 hours post treatment with doxorubicin respectively, stained with annexin V (Annexin V-FITC conjugate; Invitrogen, CO; #V13242) and propidium iodide (PI; Invitrogen, CO; #V13242), and analyzed by flow cytometry using the BD FACS Canto II cytometer (BD Biosciences, USA). Briefly, cells were harvested after incubation period and washed twice by centrifugation (1,200 g, 5 min) in cold phosphate-buffered saline (DPBS; Gibco, CO; #14040133). After washing, cells were resuspended in 0.15 ml AnnV binding buffer 1X (ABB 5X, Invitrogen, CO; #V13242; 50 mM HEPES, 700 mM NaCl, and 12.5 mM CaCl2 at pH 7.4) containing fluorochrome-conjugated AnnV and PI (PI to a final concentration of 1 ug/ml) and incubated in darkness at room temperature for 15 min. As soon as possible cells were analyzed by flow cytometry, measuring the fluorescence emission at 530 nm and >575 nm. Data were analysed using the FlowJo software version 10.5.3 (https://www.flowjo.com).

## Declarations

### Ethics approval and consent to participate

Not applicable

### Consent for publication

Not applicable

### Availability of data and materials

The dataset supporting the conclusions of this article is included as **Additional Files 1-5**. A web portal using the Shiny framework in R has been developed to visualize rank profiles of the DRIVE shRNA screen and corresponding gene expression data from TCGA at https://apsic.shinyapps.io/APSiC/. The code for the APSiC algorithm is available at https://github.com/hesmon/APSiC/. The raw shRNA data has already published as a part of the project DRIVE (https://data.mendeley.com/datasets/y3ds55n88r/4) and copy number and mutation profiles of the cell lines are available at the Cancer Cell Line Encyclopedia portal (https://portals.broadinstitute.org/ccle/home). The gene expression data from the TCGA are available at the TCGA Genomics Data Commons data portal (https://portal.gdc.cancer.gov/).

### Authors’ contributions

H.M., S.P. and C.K.Y.N. conceived the study. S.P. and C.K.Y.N. supervised the study. H.M. and C.K.Y.N. developed the methodology. H.M. performed the bioinformatic analyses. E.Z. contributed to data presentation and visualization. H.M. and E.Z. implemented the Shiny web application. M.C.-L., G.B., S.T.-M., V.P. and M.L. performed *in vitro* experiments. S.S., A.dW., G.R., M.B., N.B., M.vF. and L.M.T. critically discussed the results. H.M., M.C.-L., G.B., S.P. and C.K.Y.N. interpreted the results and wrote the manuscript. All authors agreed to the final version of the manuscript.

## Acknowledgments

Development of APSiC was performed at sciCORE scientific computing center at the University of Basel and at the scientific computing center of the Department of Bioinformatics, University of Tehran.

## Additional files

**Additional file 1:** APSiC analysis of mutation oncogenes in DRIVE

**Additional file 2:** APSiC analysis of amplification oncogenes in DRIVE

**Additional file 3:** APSiC analysis of mutation tumor suppressors in DRIVE

**Additional file 4:** APSiC analysis of non-genetic oncogenes in DRIVE

**Additional file 5:** APSiC analysis of non-genetic tumor suppressors in DRIVE

**Additional file 6:** Significant genetic drivers identified by APSiC in the pan-cancer analysis of DRIVE

**Additional file 7:** Rank profiles of *DDX27*, *DCAF8L2* and *RBM39* in DRIVE

**Additional file 8:** Pathological annotation in the DRIVE project and the corresponding TCGA projects used for gene expression analysis.

**Additional file 8**: **a**, (left) Rank profile of the *LRRC4B* gene in breast cancer cell lines. Each bar in the waterfall plots represents one cell line and is colored by the mutation status. (right) *LRRC4B* transcript expression in breast cancers and normal tissues. The plot was generated using gene expression data obtained from the TCGA dataset. **b**, Screening of LRRC4B protein expression in a panel of breast cancer cell lines by western blot. Ratio of LRRC4B expression relative to actin were calculated for each cell line. Ratio are shown above the western blot plot. **c**, Dot plot showing the flow cytometry gating strategy used to assess cell cycle status on breast cancer cell lines in both overexpressing and downregulating *LRRC4B* cells. (*** *P* < 0.001).

