## Additional File 6 for "Systematic Identification of Novel Cancer Genes through Analysis of Deep shRNA Perturbation Screens"

**Additional file 2:** Significant genetic drivers identified by APSiC in the pan-cancer analysis of DRIVE

| | Gene | Predicted function | Known function | Cancer type | Mechanisms of action |
|---|---|---|---|---|---|
| 1 | *KRAS* | Mutation/ amplification oncogene | Oncogene | Gastric cancer, squamous cell carcinoma, hepatic angiosarcoma, thyroid cancer, and others | MAPK and PI3K-AKT pathways, cell proliferation and survival, metabolism |
| 2 | *TP53* | Mutation oncogene | Oncogene/ Tumor suppressor | Leukemia, bladder cancer, breast cancer, melanoma, squamous cell carcinoma, liver cancer, kidney cancer, and others | DNA repair, cell-cycle arrest, apoptosis, senescence, autophagy, angiogenesis |
| 3 | *BRAF* | Mutation oncogene | Oncogene | Thyroid cancer, melanoma, and others | MAPK pathway, cell proliferation and survival, differentiation |
| 4 | *CTNNB1* | Mutation oncogene | Oncogene | Gastric cancer, liver cancer, kidney cancer | WNT pathway, cell proliferation and survival, epithelial-mesenchymal transition |
| 5 | *NRAS* | Mutation oncogene | Oncogene | Gastric cancer, melanoma, angiosarcoma | MAPK and PI3K-AKT pathways, cell proliferation and survival, metabolism, autophagy |
| 6 | *PIK3CA* | Mutation oncogene | Oncogene | Ovarian cancer, breast cancer, liver cancer, and others | PI3K-AKT pathway, cell proliferation and survival, metabolism, autophagy |
| 7 | *DDX27* | Mutation oncogene | Oncogene | Colorectal cancer[1], gastric cancer[2] | NF-κB pathway, cell proliferation, migration, metastasis |
| 8 | *DCAF8L2* | Mutation oncogene | NA | NA | NA |
| 9 | *RBM39* | Mutation oncogene | Oncogene | Leukemia[3,4] | RNA binding protein splicing network in AML, anticancer drug (splicing inhibitors) resistance[5], mTOR signalling pathway[5,6] |
| 10 | *CDK4* | Amplification oncogene | Oncogene | Non-small cell lung cancer[7,8], cervical cancer[7], melanoma[7,9], liposarcoma[10] and others | Cell-cycle check-point regulation, metabolism (anaerobic glycolysis)[11] |
| 11 | *TLK2* | Amplification oncogene | Oncogene | Breast cancer[12,13], glioblastoma[14] | Genomic instability[12], src signalling pathway[14] |
| 12 | *IL6* | Amplification oncogene | Oncogene | Liver cancer[15], breast cancer[16], head and neck squamous cell carcinoma[17], prostate cancer[15,18] | STAT3 activation, insulin-growth factor signaling, inflammation |
| 13 | *HAS2* | Amplification oncogene | Putative oncogene | Breast cancer[19] | Radio-resistance[19,20], epithelial-mesenchymal transition[21], cell proliferation, apoptosis[21,22] |
| 14 | *YAP1* | Amplificatio | Oncogene | Epithelial cancers (colorectal, | Hippo, Notch, PI3K-mTOR, MEK, |

| | | | | | |
|---|---|---|---|---|---|
| | | n oncogene | | pancreatic, gastric, liver, breast, NSCLC) | Wnt/β-catenin pathways, stemness |
| 15 | *ARID1A* | Mutation tumor suppressor | Tumor suppressor | Liver cancer, gastric cancer, colorectal cancer, ovarian and endometrial cancers | Chromatin remodelling, epigenetic regulation, PI3K/AKT pathway activation |
