## Supplementary figures and images for "Systematic Identification of Novel Cancer Genes through Analysis of Deep shRNA Perturbation Screens"

### Additional File 7

#### Additional file 3

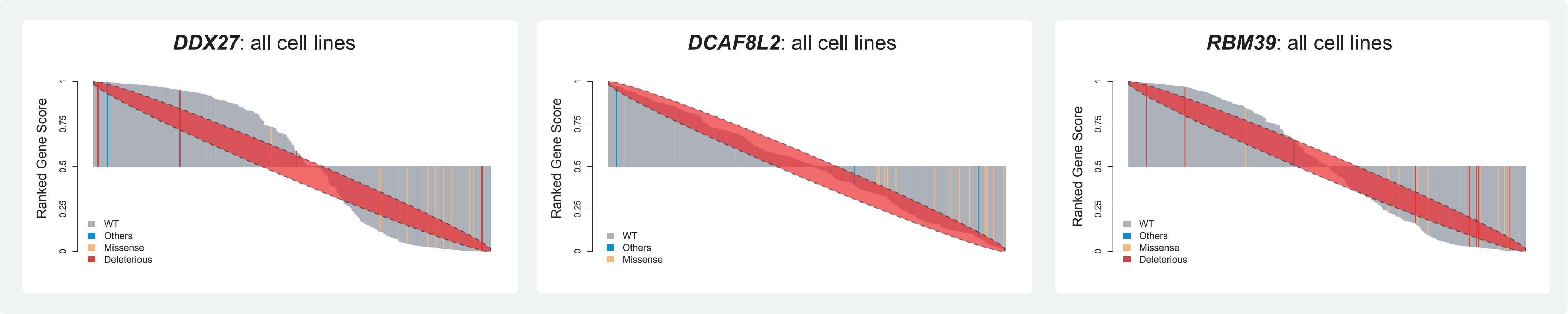

### Additional File 9

#### Additional file 5

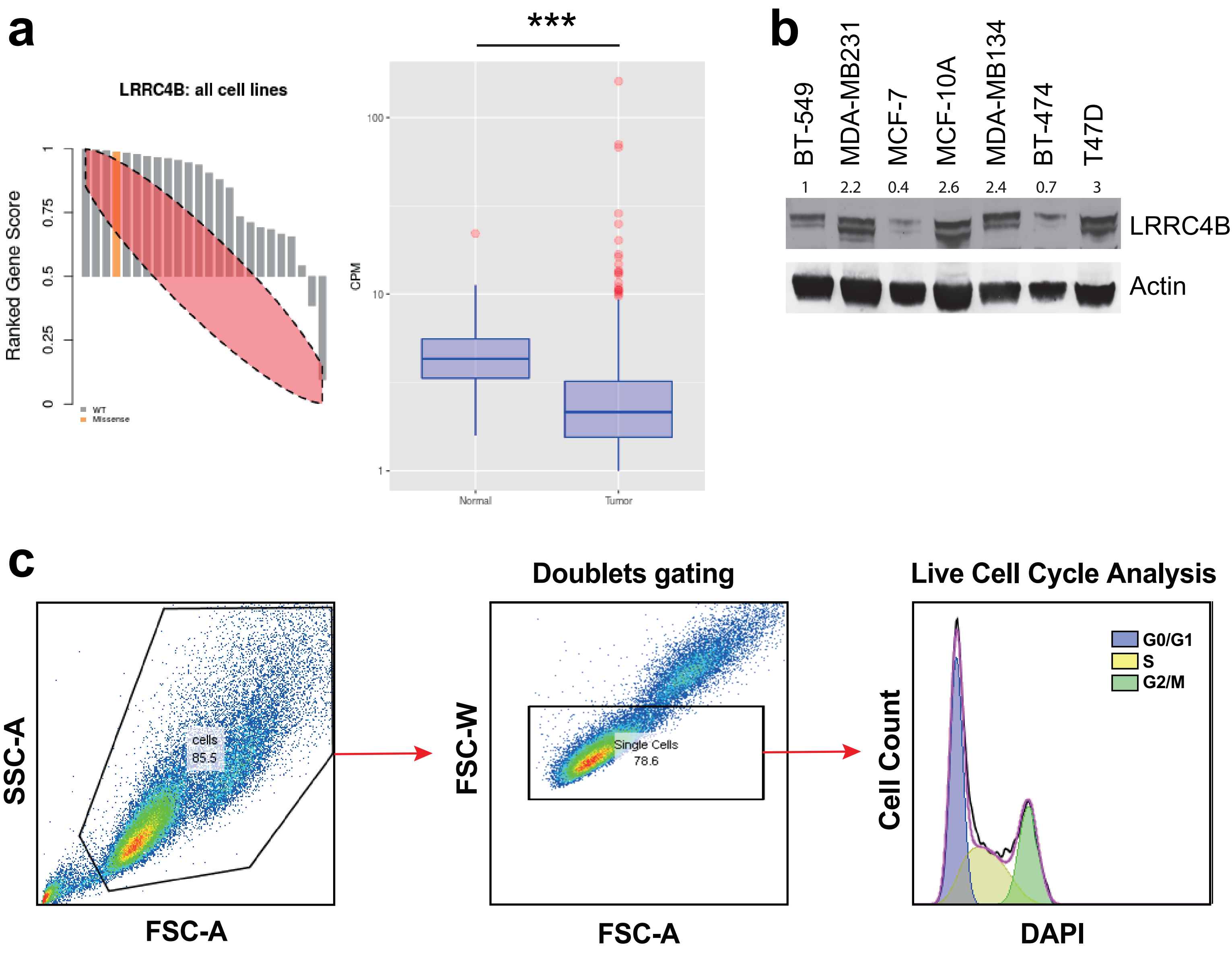
