## Additional File 8 for "Systematic Identification of Novel Cancer Genes through Analysis of Deep shRNA Perturbation Screens"

### Additional file 4

| Pathological annotation in the DRIVE project | TCGA project | Pathological annotation in the DRIVE project | TCGA project |
|---|---|---|---|
| Oesophagus Carcinoma | ESCA | Bladder Carcinoma | BLCA |
| Upper Aerodigestive Tract Carcinoma | HNSC | Liver_HCC | LIHC |
| Lung_NSCLC Squamous | LUSC | Thyroid Carcinoma | THCA |
| Colorectal Carcinoma | COAD, READ | Kidney Carcinoma | KIRC, KIRP |
| Gastric Carcinoma | STAD | Lung_NSCLC Adenoma | LUAD |
| Breast Carcinoma | BRCA | Endometrium Carcinoma | UCEC |
| Lung_NSCLC Large_Cell, Leukemia_AML, Leukemia_ALL, Lymphoma NH_B_cell, Soft_Tissue Sarcoma Rhabdoid, PNET Neuroblastoma Skin_Melanoma, Ovary Carcinoma, CNS_Glioma, CNS_Glioma HighGrade, Lung_NSCLC Others, Lung_SCLC, Pancreas Carcinoma, Lung Mesothelioma | | | Not Available |
